# Linoleic acid supplementation reverses microglial response to diet induced-obesity at hypothalamic, cortical and subcortical level in mice

**DOI:** 10.1101/2023.07.21.549983

**Authors:** Lucas Jantzen, Stéphanie Dumontoy, Bahrie Ramadan, Christophe Houdayer, Emmanuel Haffen, Aziz Hichami, Naim Khan, Vincent Van Waes, Lidia Cabeza

## Abstract

Obesity is a major risk factor for neuropsychiatric alterations. Fatty regimes lead to systemic and cerebral inflammation, the latest acting through lipotoxicity on hypothalamic structures controlling energy homeostasis. Since literature points to a protective effect of linoleic acid (LA) on mood disorders through the regulation of systemic inflammation, we investigated how five weeks of LA supplementation modulates emotional behaviour and microglia-related neuroinflammation. C57BL/6j mice were fed with either a high-fat (HFD) or a standard diet for 12 weeks, underwent a battery of behavioural tests and were subsequently sacrificed for immunofluorescence staining targeting microglia-specific calcium-binding proteins (IBA-1). Neuroinflammation severity was approximated in multiple hypothalamic, cortical and subcortical regions. Our results show an anxio-depressive-like effect of sustained HFD that neither was alleviated nor worsen with LA supplementation. Increased IBA-1 expression in the HFD group was substantially reversed with LA supplementation. Thus, our results suggest anti-neuroinflammatory properties of LA not restricted to hypothalamic areas, but also evident at the cortical and subcortical level. This study is therefore relevant in the frame of obesity and neuropsychiatric disorders with a neuroinflammatory basis. Further investigation may provide more information to justify dietary strategies aiming at reducing the impact of obesity associated comorbidities.

## Introduction

The multifactorial aetiology of obesity entangles the comprehension of the underlying biological mechanisms [1]. Few genetic factors increasing the vulnerability of individuals to suffer from obesity throughout their lives have been identified, as for example, a defective hypothalamic melacortin 4 receptor [for review ^2^]. However, a large fraction of obesity cases in Western countries is a consequence of malnutrition, i.e. an overconsumption of fatty food also called *junk food*. These high-fat regimes induce other metabolic alterations that in the long-run are responsible for the onset of cardiac, respiratory and endocrinal comorbidities [3,4]. Additionally, obesity has been linked to neuropsychiatric disorders as anxiety and depression [5,6], with clinical [7–9] and preclinical studies [10–13] demonstrating a positive correlation for the co-existence of these conditions.

Obesity has reached much scientific attention since Carabotti and colleagues, in 2015, demonstrated the crucial role of microbiota in human health [14]. Increased short-chain fatty acids (SCAFs) faecal concentrations have been reported in individuals suffering from obesity [15], as a consequence of changes in their microbiome. In physiological conditions, these products of colon microbial fermentation induce the release of anorexigenic hormones into the peripheral circulation [16], as well as they increase circulating concentrations of leptin and insulin [17]. Since they are able to cross the intestinal epithelium, reach the portal vein and stimulate the vagal nerf, SCAFs therefore influence the homeostatic balance controlling individuals’ appetite at short- and long-term. Additionally, the SCAFs are modulatory agents of the immune response: they can reduce peripheral inflammation by inhibiting the production of pro-inflammatory cytokines from monocytes and macrophages [18], and most importantly, they participate in the maturation of the microglia in the central nervous system (CNS), and so contribute to its defence [19]. Subsequently, abnormal high circulating SCAFs concentrations in obesity correlate to higher concentrations of lipopolysaccharides (LPS), and therefore to the severity of the consequent systemic inflammation [20]. Additionally, the expression of gustatory receptors, which is sensitive to LPS circulating levels, is downregulated in obese individuals, which leads to an altered gustatory perception and may motivate food overconsumption [for review_21]._

From the time it was first described in the hypothalamus [22], obesity-induced neuroinflammation has garnered significant scientific attention. Different mechanisms have been described in an attempt to unravel the role of neuroinflammation in the pathogenesis of obesity, and aiming to develop new therapeutic strategies against body weight gain. An altered blood-brain barrier integrity in terms of both structure and function, will admit a dynamic passage of molecules from the vascular system, including immune cell infiltration [23–26]. Besides, dietary regimes enriched in fatty acids are known to upregulate nuclear factor kappa-B (NF-κB) inflammatory signalling and therefore to modulate energy homeostasis [27], particularly in hypothalamic regions. In this context, and since dietary components act as modulatory actors of the inflammatory pathways in the CNS, investigation on the impact of supplemented regimes targeting bioactive metabolites with anti-inflammatory potential stands to reason.

Linoleic acid (LA) is an essential omega-6 (ω-6) polyunsaturated fatty acid present in vegetables oils, nuts, seeds, meats and eggs [28]. As other ω-6 fatty acids, LA participates in the maintenance of the cellular membrane structure, and mainly throughout its metabolite gamma linoleic acid, influence its function. It also participates in the preservation of nerve blood flow through the formation of prostaglandin, and as a precursor of eicosanoids, it modulates vital functions (e.g. pulmonary function), vascular tone and the inflammatory response [for review ^29^]. Linoleic acid has become ubiquitous in Western diets due to the incorporation of high-LA soybean and corn oils, and represent approximately 7% of the intake of daily calories [for review ^28^]. Thus, the pathogenicity of high-fat regimes seems to depend more on the ratio between ω-6 and ω-3 fatty acids (e.g. alpha-linolenic acid) [30], and in relation to the present fraction of saturated fatty acids.

Preclinical studies in murine models have demonstrated that hypothalamic microglial-related inflammatory response induced by short-term high-fat regimes can be rescued by linoleic acid [31]. However, neuroinflammation chronicity might limit the mechanisms from which LA exerts its anti-inflammatory effect. We therefore propose to evaluate overall brain effects of LA supplementation on obesity-related neuroinflammation induced through a sustained highly saturated fat regime as well as its effects on emotional balance. We expect to confirm the emergence of an anxio-depressive-like phenotype consequent to the sustained rich-fat regime, and hypothesize that LA anti-neuroinflammatory effects will also alleviate these emotional disturbances.

## Results

The timeline of the experimental design is presented in **Figure 1a**.

**Figure 1.**
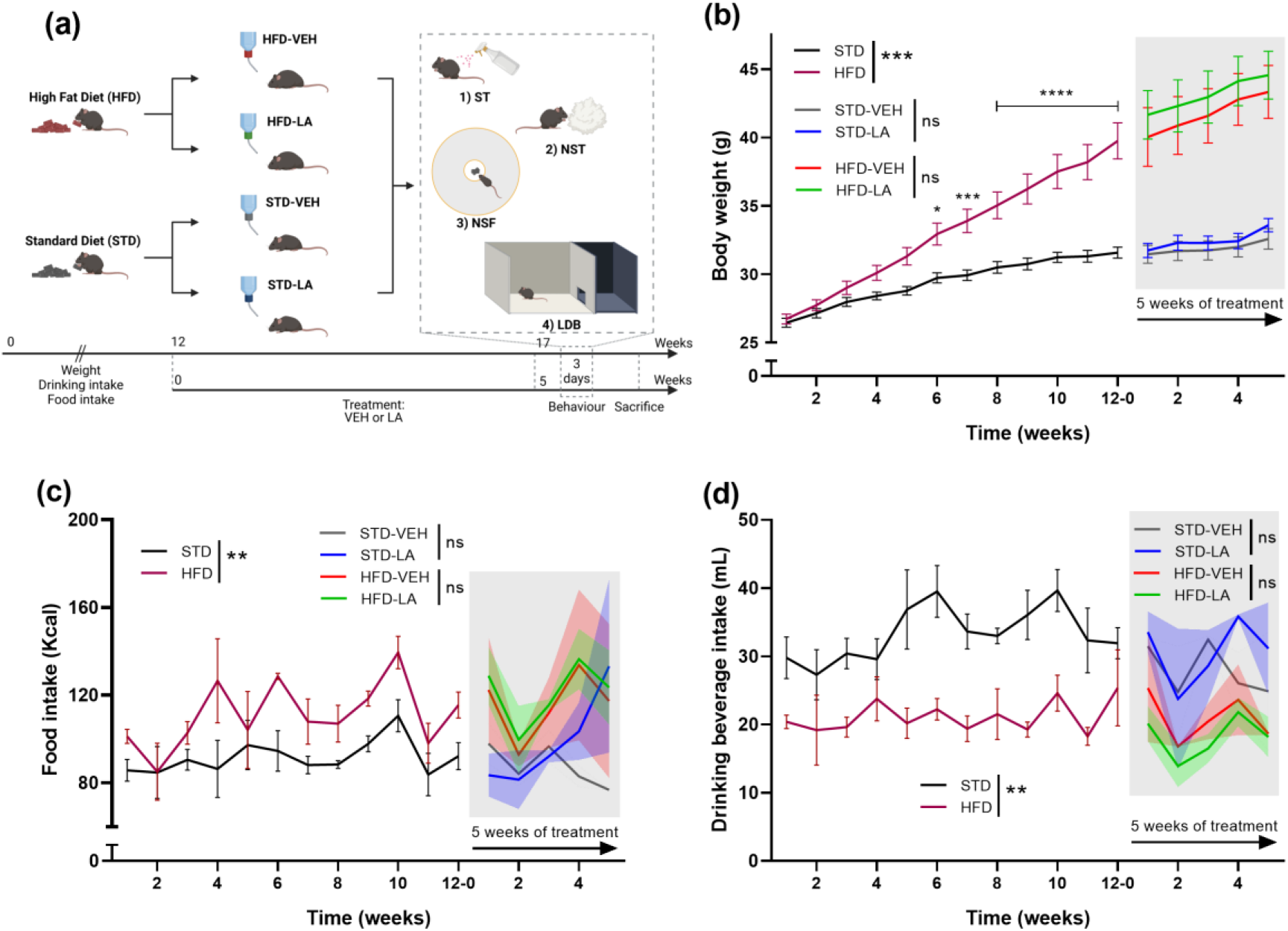
Experimental design and animals’ monitoring. **(a)** Animals were fed with either a high-fat (HFD) or a standard (STD) diet for 12 weeks before starting being treated with vehicle (VEH) or a 0.2% linoleic acid (LA) solution during five weeks. The body weight of animals and their consummatory behaviour, in terms of food and drinking beverage intake, was monitored during the entire duration of the experiment (3 times per week). The effect of the regime and of LA supplementation on emotional behaviour was evaluated during the last week of the experiment through a battery of tests: the splash test (ST), the nestlet shredding test (NST), the novelty supressed feeding (NSF) task and the light-dark box (LDB) test. Animals were subsequently sacrificed for IBA-1 immunofluorescence staining. Created with *BioRender.com*. **(b)** All animals progressively gained weight during the experiment, and mice fed with HFD weighted significantly more than their control counterparts (STD) from the sixth week of differential regime. However, five weeks of LA supplementation did not affect the body weight of animals, neither for the HFD nor for the STD group. **(c)** A general effect of the regime on the weekly food intake was evidenced, with HFD mice consuming significantly more calories than control animals. Besides, LA supplementation did not significantly impact food intake neither in obese (HFD-LA vs HFD-VEH) nor in non-obese individuals (STD-LA vs STD-VEH). **(d)** Sustained fat intake significantly influenced the weekly drinking intake of mice, with HFD individuals consuming a significantly lower volume of drinking beverage than control animals. However, LA supplementation did not modify this behaviour neither in obese (HFD-LA vs HFD-VEH) nor in non-obese individuals (STD-LA vs STD-VEH). Values plotted are mean ± SEM (n=10 per group). *, p<0.05; **, p<0.01; ***, p<0.001; ****, p<0.0001; ns, not significant.

### Body weight and consummatory behaviour

The body weight of animals, and their food and drinking beverage intake, were monitored three times a week and revealed a significant increase of the body mass in those maintained under a high-fat diet (-HFD; mean weight at week 12^th^ (g) ± SEM: 39.75±1.32) compared to mice fed with standard food (-STD; 31.59±0.40) [RMA: main effect of the diet, F_1,38_=18.0, p<0.001; mean effect of time, F_11,418_=178.9, p<0.0001; diet x time interaction, F_11,418_=34.0, p<0.0001] (**Figure 1b**). As expected, animals from the HFD group consumed significantly more calories than control animals (mean food intake per week (Kcal) ± SEM, HFD: 111.30±4.36; STD: 91.72±2.20) [diet: F_1,6_=24.2, p<0.01; time: F_11,66_=1.6, p<0.01; diet x time interaction, F_11,66_=0.7, p=0.70], and their drinking intake was lower (mean volume of beverage intake per week (mL) ± SEM, HFD: 21.15±2.35; STD: 33.33±3.96) [diet: F_1,6_=23.6, p<0.01; time: F_11,66_=1.6, p=0.11; diet x time interaction, F_11,66_=1.1, p=0.41]. This consummatory behaviour did not change during the period of LA supplementation (food intake, HFD-VEH: 115.70±6.74; HFD-LA: 120.80±6.29; STD-VEH: 87.72±4.12; STD-LA: 98.79±9.45; treatment intake, HFD-VEH: 20.97±1.57; HFD-LA: 18.10±1.38; STD-VEH: 27.93±1.67; STD-LA: 30.57±2.09) [food: F_3,4_=1.8, p=0.29; time: F_4,16_=1.6, p=0.21; food x time interaction, F_12,16_=0.8, p=0.66; treatment: F_3,4_=2.6, p=0.19; time: F_4,16_=3.1, p<0.05; treatment x time interaction, F_12,16_=0.5, p=0.88] (**Figure 1c,d**).

### Behavioural evaluation

The battery of behavioural tests used in this study allowed confirming the emergence of an anxio-depressive-like phenotype in mice as a consequence of a prolonged rich-fat regime, in agreement with the literature[10,11,32]. Compared to STD, sixteen-to-seventeen weeks of HFD significantly reduced the time mice spent grooming themselves after being exposed to a viscous 15% sucrose solution (mean of the total grooming time (s) ± SEM; STD-VEH: 61.90±7.29; HFD-VEH: 43.40±6.42) [KW, H_3,40_=13.2, p<0.01; *post-hoc* MWU, Z=2.05, p<0.05] (**Figure 2a**). However, animals of both conditions started grooming at similar latencies (mean of the grooming latency (s) ± SEM; STD-VEH: 81.22±10.49; HFD-VEH: 72.23±5.63) [H_3,40_=4.3, p=0.23; Z=0.076, p=0.94] (**Figure 2b**). Contrary to expected, HFD-VEH and STD-VEH mice behaved similarly in the NSF task, displaying comparable latencies to approach the available food (mean latency (s) ± SEM; STD-VEH: 25.10±4.79; HFD-VEH: 32.80±7.74) [GBW: Χ^2^=0.57, p=0.451] (**Figure 2c**). Furthermore, HFD-VEH animals displayed a typical anxious behaviour in the LDB test, spending significantly less time in the light compartment (mean of the time in the light compartment (s) ± SEM; HFD-VEH: 197.15±14.03) than their SDT-VEH counterparts (278.67±14.20) [H_3,40_=20.0, p<0.000; Z=3.10, p<0.01] (**Figure 2d**). However, no effect of the food regime was evidenced in terms of the frequency of visiting the light compartment (mean of entries in the light compartment ± SEM; HFD-VEH: 26.00±3.57; STD-VEH: 37.70±4.84) [H_3,40_=6.0, p=0.11] (**Figure 2e**), the time spent in the transition zone (mean of time spent in the transition zone (s) ± SEM; HFD-VEH: 69.67±9.72; STD-VEH: 82.81±8.26) [H_3,40_=1.7, p=0.63] (**Figure 2f**) and the total distance travelled during the test (mean of the travelled distance (cm) ± SEM: STD-VEH: 3494.12±188.37; HFD-VEH: 3096.50±207.08) [H_3,40_=3.6, p=0.30] (**Figure 2g**). From the evaluation of the effect of sustained rich-fat regime on promoting the emergence of compulsive behaviours through the NS test, no differences were found between the experimental conditions (mean of shredded material (%) ± SEM; STD-VEH: 2.04±0.34; HFD-VEH: 1.87±0.46) [H_3,40_=3.3, p=0.34] (**Figure 2h**).

**Figure 2.**
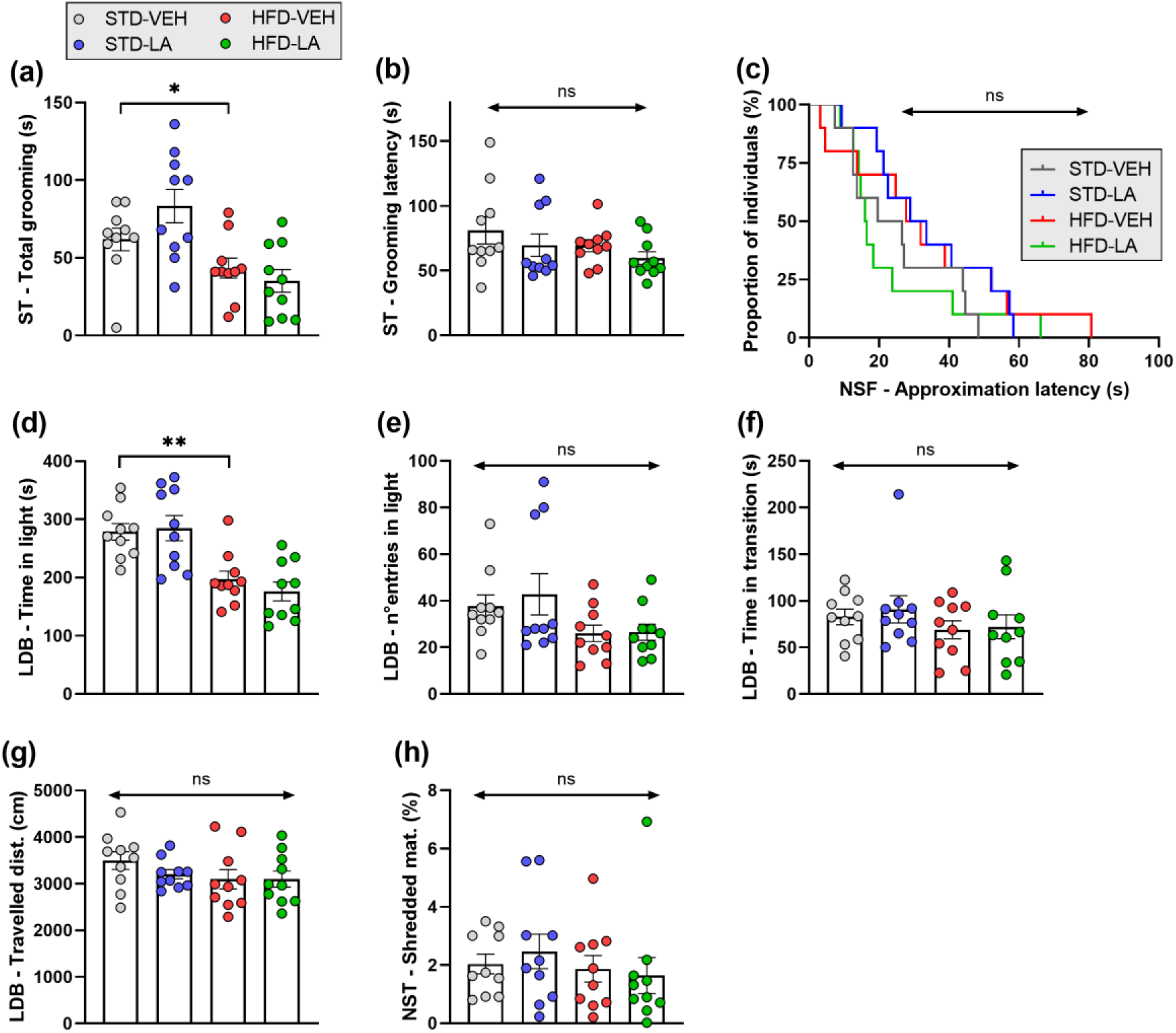
Effect of five weeks of linoleic acid (LA) supplementation on sustained high-fat diet (HDF)-induced emotional alterations. **(a)** A depressive-like phenotype consequent to sustained HFD was evidenced by the splash test (ST), with obese individuals (HFD-VEH) spending less time grooming themselves than individuals fed with a standard diet (STD-VEH). **(b)** However, the latency of the first grooming event in the ST was similar for both experimental groups. **(a,b)** Linoleic acid supplementation did not affect neither the behaviour of obese (HFD-LA) nor that of non-obese individuals (STD-LA) in the ST. **(c)** The behavioural evaluation through the novelty suppressed feeding (NSF) task did not reveal emotional differences between the experimental groups, with all animals requiring similar latencies to approach the available food. **(d-g)** The light-dark box (LDB) test allowed revealing the emergence of an anxious-like phenotype consequent to a sustained HFD, with obese individuals (HFD-VEH) spending less time in the light compartment that their control counterparts (STD-VEH) **(d)**. However, this emotional alteration was no revealed through the other parameters measured, i.e. frequency of visiting the light compartment **(e)**, time in the transition area **(f)** and travelled distance **(g)**. Besides, no behavioural effect of five weeks of LA supplementation was evidenced in the LDB test **(d-g)**, irrespective of the animals’ regime. **(h)** Sustained HFD did not induce the emergence of repetitive, compulsive-like behaviours in mice as evaluated in the nestlet shredding test (NST) and no effect of LA supplementation was observed, irrespective of the animals’ food regime. Values plotted are mean ± SEM (n=10 per group). *, p<0.05; **, p<0.01; ns, not significant.

The effect of 5 weeks of supplemented LA on animals’ behaviour was subsequently studied. Contrary to expected, no differences between the experimental groups were found concerning the anxio-depressive- and compulsive-like behaviours evaluated, neither for HFD mice (HFD-LA: SP, grooming time: 35.10±7.28; grooming latency: 59.78±4.88; NSF, approach latency: 22.80±5.58; LDB, time in light: 176.06±15.94; entries in light: 26.50±3.43; time in transition: 72.28±12.77; travelled distance: 3098.45±172.83; NS, shredded material: 1.64±0.62, [HFD-VEH vs HFD-LA: Χ^2^=0.831, and all ps>0.1], nor for their STD counterparts (STD-LA: SP, grooming time: 83.30±10.77; grooming latency: 69.67±8.73; NSF, approach latency: 33.90±5.44; LDB, time in light: 284.98±21.67; entries in light: 42.80±8.81; time in transition: 90.88±14.59; travelled distance: 3204.34±98.62; NS, shredded material: 2.47±0.60 [STD-VEH vs STD-LA: Χ^2^=1.46, and all ps>0.1] (**Figures 2a-h**).

### IBA-1 expression

Following the behavioural evaluation and sacrifice of animals, IBA-1 relative expression as a proxy of microglial response, was assessed through immunofluorescence. The values obtained for each selected brain region are detailed in **Table 1**. Values are relative to the control group STD-VEH (mean = 100%).

**Table 1.** Data of IBA-1 fluorescence intensity relative to the control group (standard diet-vehicle – STD-VEH; 100%) for the experimental conditions and the brain regions of interest (ROI). Food regime: high-fat diet –HFD. Treatment: linoleic acid supplementation –LA. Data are means ± SEM. Paraventricular nucleus of the hypothalamus –PVN, peduncular part of the lateral hypothalamus – PLH, ventromedial hypothalamus –VMH, lateral hypothalamus –LH, subthalamic nucleus of the hypothalamus –STN, parasubthalamic nucleus of the hypothalamyus PSTN, prelimbic cortex -PL, infralimbic cortex –IL, orbitofrontal cortex –OFC, medial anterior cingulate cortex -ACC, anterior insular cortex -aIC, dorsomedial striatul –DMS, dorsolateral striatum –DLS, dorsal region of the bed nucleus of the stria terminalis –dBNST, dorsal hippocampus -dHipp, lateral habenula –LHb, basolateral amygdala –BLA, central amygdala –CeA, ventral posteromedial thalamic nucleus -VPM, and parvicellular part of the ventral posterior thalamic nucleus -VPPC.

| ROI | IBA-1 relative fluorescence intensity (mean % ± SEM) | | |
| --- | --- | --- | --- |
|  | STD-LA | HFD-VEH | HFD-LA |
| *Hypothalamic regions* |  |  |  |
| *PVN* | 111.50±11.84 | 149.50±11.35 | 108.20±8.04 |
| *PLH* | 116.20±11.71 | 140.90±8.01 | 105.80±7.68 |
| *VMH* | 103.10±8.09 | 130.70±10.45 | 114.50±9.26 |
| *LH* | 116.00±8.89 | 131.00±13.61 | 106.80±9.53 |
| *STN* | 152.50±3.04 | 159.80±24.60 | 122.60±10.72 |
| *PSTN* | 161.10±7.98 | 158.60±17.60 | 109.50±9.21 |
| *Cortical regions* |  |  |  |
| *PL* | 110.20±9.52 | 140.00±9.47 | 99.83±6.36 |
| *IL* | 104.30±7.97 | 140.90±12.30 | 104.20±6.64 |
| *OFC* | 95.65±7.86 | 143.01±3.63 | 99.55±7.55 |
| *ACC* | 92.29±7.59 | 136.10±12.56 | 99.93±6.43 |
| *aIC* | 115.90±8.44 | 141.40±6.41 | 109.50±10.17 |
| *Subcortical regions* |  |  |  |
| *DMS* | 90.00±5.13 | 151.50±16.59 | 105.30±3.26 |
| *DLS* | 104.30±3.88 | 159.70±13.65 | 107.90±7.30 |
| *dBNST* | 89.15±6.57 | 153.20±10.71 | 103.90±5.62 |
| *dHipp* | 90.07±5.24 | 139.80±17.11 | 100.20±8.18 |
| *LHb* | 121.70±8.34 | 142.50±13.38 | 104.40±8.75 |
| *BLA* | 122.10±6.34 | 144.10±13.19 | 103.40±9.86 |
| *CeA* | 138.80±10.91 | 147.70±17.72 | 99.42±14.81 |
| *VPM* | 121.90±3.07 | 201.70±38.52 | 142.10±18.63 |
| *VPPC* | 115.80±7.62 | 140.20±16.46 | 86.53±6.19 |

In order to confirm the neuroinflammatory effect of sustained rich-fat regimes already described in the literature, IBA-1 expression was first evaluated in six hypothalamic brain structures (in rostral-to-caudal order: paraventricular nucleus -PVN, ventromedial hypothalamus -VMH, peduncular part of the lateral hypothalamus -PLH, lateral hypothalamus -LH, subthalamic nucleus -STN and parasubthalamic nucleus -PSTN). A main effect of HFD was evidenced in the PVN [KW, H_3,29_=10.0, p<0.05; *post-hoc* MWU, Z=-2.8, p<0.01], the PLH [H_3,29_=11.2, p<0.05; Z=-2.9, p<0.01] and the PSTN [H_3,27_=16.0, p=0.001; Z=-2.8, p<0.01], as well as a tendency in the STN [H_3,29_=10.0, p<0.05; Z=-1.8, p=0.07]. However, no differential expression of IBA-1 between experimental conditions was found in the VMH [H_3,29_=5.9, p=0.12], nor in the LH [H_3,29_=2.7, p=0.45] (**Figure 3**). Subsequently, the effect of the regime was evaluated in various cortical structures (in rostral-to-caudal order: prelimbic cortex - PL, infralimbic cortex –IL, orbitofrontal cortex –OFC, medial anterior cingulate cortex -ACC, anterior insular cortex -aIC), showing a generalized significant increase of IBA-1 expression with the only exception of the IL (PL [H_3,29_=7.9, p<0.05; Z=-2.5, p<0.05], IL [H_3,29_=6.4, p=0.09], OFC [H_3,29_=17.4, p=0.000; Z=-3.4, p<0.001], ACC [H_3,28_=8.4, p<0.05; Z=-2.2, p<0.05] and aIC [H_3,29_=11.9, p<0.01; Z=-3.2, p<0.01]) (**Figure 4**). Finally, the effect of sustained rich-fat regime on IBA-1 expression was studied in several sub-cortical brain regions (dorsomedial striatul –DMS, dorsolateral striatum –DLS, dorsal region of the bed nucleus of the stria terminalis –dBNST, dorsal hippocampus -dHipp, lateral habenula –LHb, basolateral amygdala –BLA, central amygdala –CeA, ventral posteromedial thalamic nucleus - VPM, and parvicellular part of the ventral posterior thalamic nucleus -VPPC), revealing a significant increase in the DMS [H_3,28_=12.5, p<0.01; Z=-2.5, p<0.05], DLS [H_3,28_=14.1, p<0.01; Z=-3.1, p<0.01], dBNST [H_3,29_=17.5, p=0.000; Z=-3.3, p<0.01], BLA [H_3,29_=8.5, p<0.05; Z=-2.2, p<0.05], CeA [H_3,29_=8.9, p<0.05; Z=-2.0, p<0.05] and LHb [H_3,29_=9.2, p<0.05; Z=-2.4, p<0.05]. A tendency was also detected for VPPC [H_3,29_=9.3, p<0.05; Z=-1.9, p=0.058], and no main effect of the food regime on IBA-1 expression was evidenced in dHipp [H_3,29_=5.6, p=0.14] and VPM [H_3,29_=6.5, p=0.09] (**Figure 5**).

**Figure 3.**
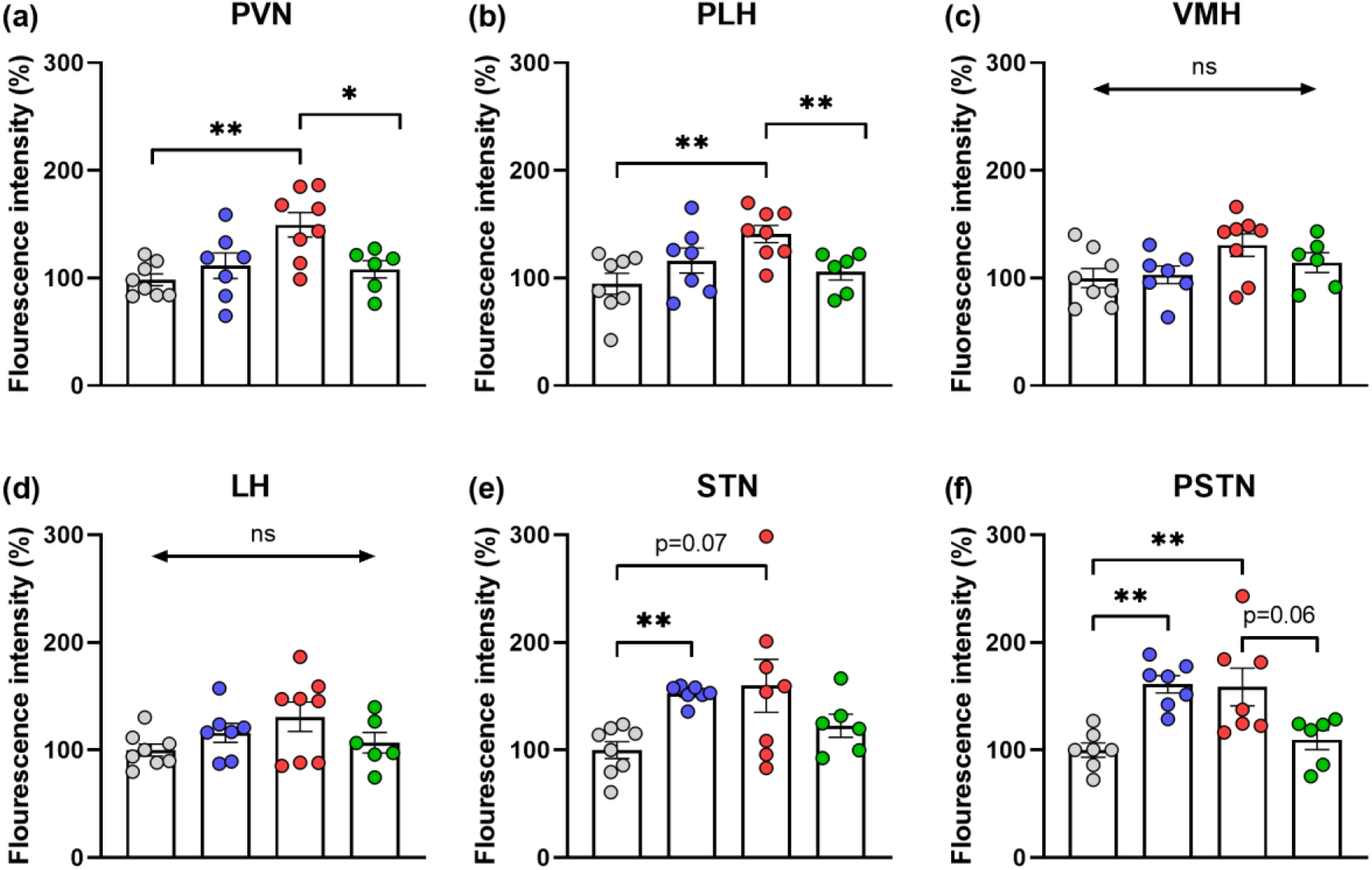
Hypothalamic IBA-1 relative expression assessed through immunofluorescence staining: evaluation of five weeks of linoleic acid (LA) supplementation on sustained high-fat diet (HDF)-induced neuroinflammation. A significant increase in IBA-1 fluorescence intensity in obese individuals (HFD-VEH) compared to control mice (STD-VEH) indicates a microgliosis consequent to the sustained rich-fat regime in various hypothalamic areas involved in the homeostatic control of food intake: paraventricular nucleus (PVN) **(a)**, penduncular part of the lateral hypothalamus (PLH) **(b)**, and parasubthalamic nucleus (PSTN) **(f)**. However, no effect of sustained HFD was observed in the ventromedial (VMH) **(c)** and in the lateral hypothalamus (LH) **(d)**, and only a tendency in the subthalamic nucleus (STN) **(e).** Supplementation of LA during five weeks significantly reduced microglial IBA-1 expression in the PVN **(a)** and PLH **(b)**, and a tendency was observed in the PSTN **(f)**. Besides, LA supplementation significantly increased IBA-1 relative expression in the STN **(e)** and PSTN **(f)** of non-obese individuals (STD-LA), indicating a special sensibility of these 2 regions to the presence of the ω-6 fatty acid in physiological conditions. No other effects on microglia-related neuroinflammation were observed. Values plotted are mean ± SEM (n=6-8 per group). *, p<0.05; **, p<0.01; ns, not significant. Colour code: grey for STD-VEH, blue for STD-LA, red for HFD-VEH and green for HFD-LA.

**Figure 4.**
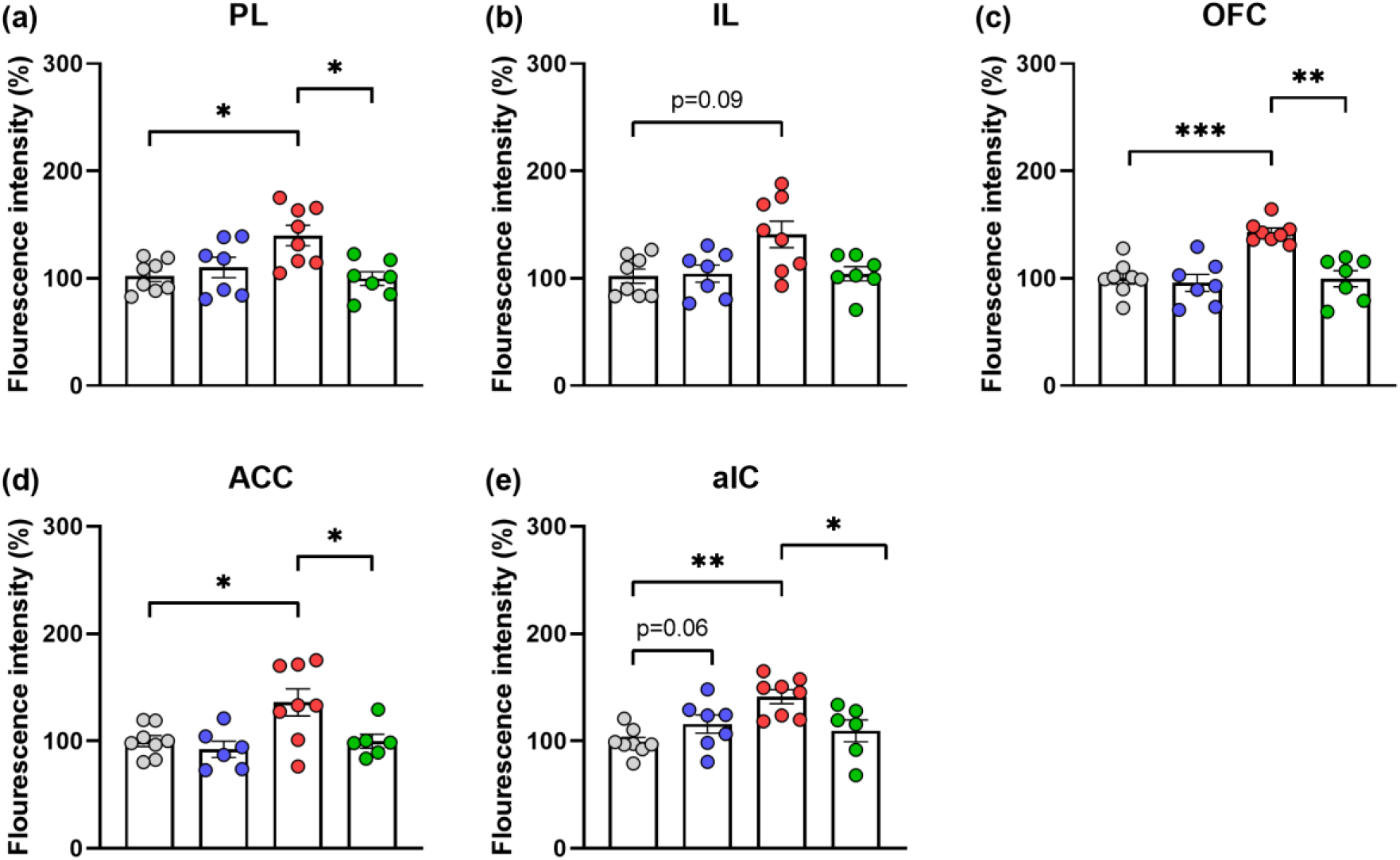
Cortical IBA-1 relative expression assessed through immunofluorescence staining: evaluation of five weeks of linoleic acid (LA) supplementation on sustained high-fat diet (HDF)-induced neuroinflammation. Significant increase of microglia-related IBA-1 fluorescence intensity in obese mice (HFD-VEH) compared to non-obese individuals (STD-VEH) in various cortical regions: prelimbic cortex (PL) **(a)**, orbitofrontal cortex (OFC) **(c)**, anterior cingulate cortex (ACC) **(d)** and anterior insular cortex (aIC) **(e)**. A tendency was also observed in the infralimbic cortex (IL) **(b)**. With the exception of the IL **(b)**, supplementation of LA during five weeks significantly reduced microglial IBA-1 expression in the cortical brain regions investigated **(a, c-e)**. No effect of LA supplementation was observed in control animals (STD-LA), besides in the aIC, where relative IBA-1 expression tend to increase compared to their control counterparts (STD-VEH) **(e)**. Values plotted are mean ± SEM (n=6-8 per group). *, p<0.05; **, p<0.01; ***, p<0.001; ns, not significant. Colour code: grey for STD-VEH, blue for STD-LA, red for HFD-VEH and green for HFD-LA.

**Figure 5.**
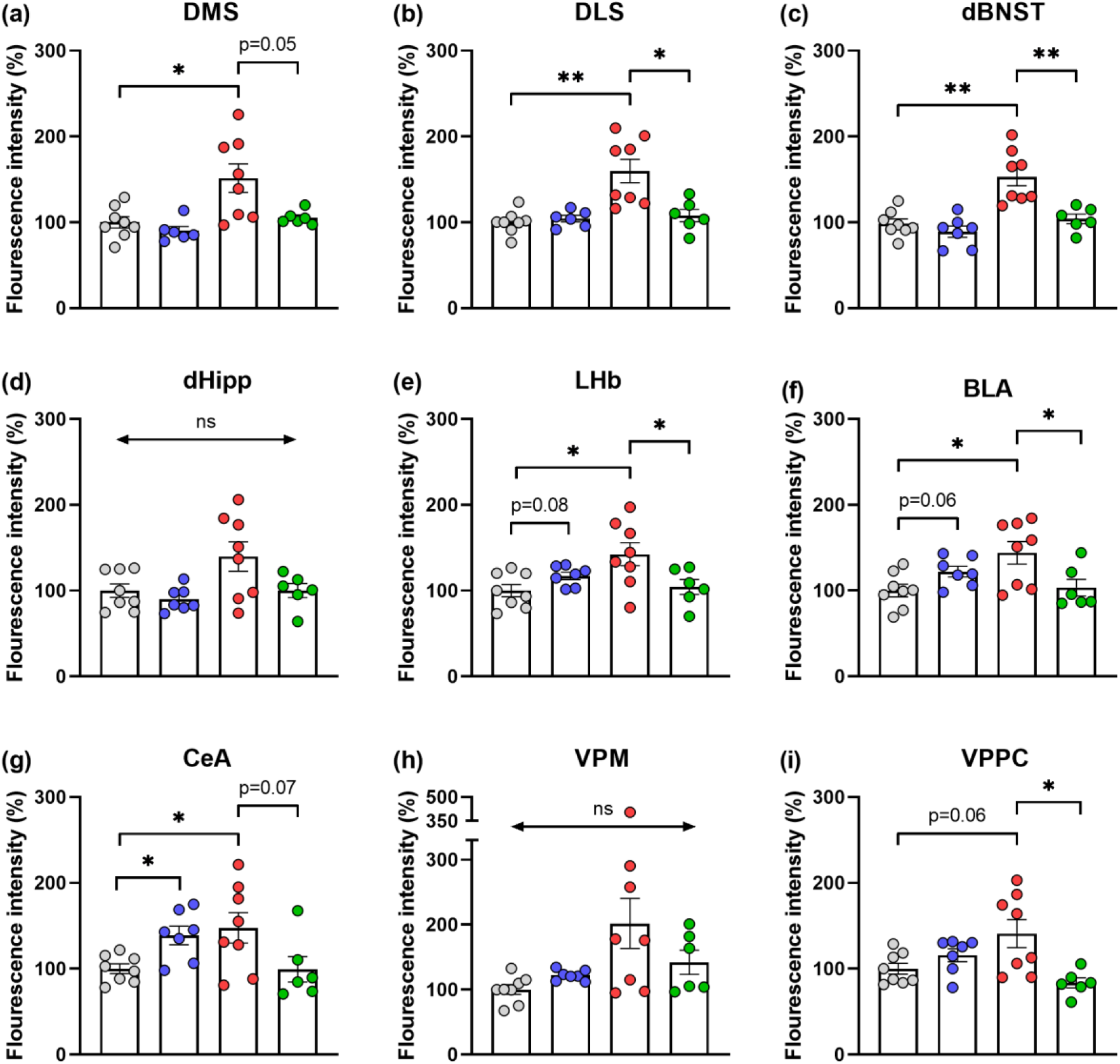
Subcortical IBA-1 relative expression assessed through immunofluorescence staining: evaluation of five weeks of linoleic acid (LA) supplementation on sustained high-fat diet (HDF)-induced neuroinflammation. Compared to animals fed with a standard regime (STD-VEH), microglia-related IBA-1 fluorescence was significantly increased in mice after 17 weeks of HFD (HFD-VEH) in various subcortical regions: dorsomedial striatum (DMS) **(a)**, dorsolateral striatum (DLS) **(b)**, dorsal part of the bed nucleus of the stria terminalis (dBNST) **(c)**, lateral habenula (LHb) **(e)**, and basolateral (BLA) **(f)** and central (CeA) **(g)** amygdala. A tendency was observed in the parvicellular part of the ventral posterior thalamic nucleus (VPPC) **(f)**. No effect of the regime on microglia-related IBA-1 relative expression was observed neither in the dorsal hippocampus (dHipp) **(d)** nor in the ventral posteromedial thalamic nucleus (VPM) **(h)**. At subcortical level, 5 weeks of LA supplementation significantly reduced microglial-related neuroinflammation in various regions **(b, c, e, f, i)**, and 2 tendencies were also observed **(a,g)**. Additionally, LA supplementation significantly increased IBA-1 relative expression of non-obese individuals (STD-LA) in the CeA **(g)**, and a tendency was observed in the LHb **(e)** and BLA **(f)**. These last results suggest a specific ω-6 fatty acid sensibility of these regions in physiological conditions. No other effects on microglia-related neuroinflammation were observed. Values plotted are mean ± SEM (n=6-8 per group). *, p<0.05; ns, not significant. Colour code: grey for STD-VEH, blue for STD-LA, red for HFD-VEH and green for HFD-LA.

Concerning the evaluation of the anti-inflammatory properties of LA supplementation in obese mice, an overall decrease of IBA-1 expression has been revealed. Hence, compared to the HFD-VEH group, HFD-LA individuals showed a reduced hypothalamic microglial response in the PVN [Z=2.2, p<0.05] and PLH [Z=2.6, p<0.01], and a tendency in the PSTN [Z=1.9, p=0.06]. However, no significant beneficial effect of LA supplementation was observed in the STN [Z=0.9, p=0.37] (no main effect observed in the VMH and LH; KW, p>0.05) (**Figure 3**). Six weeks of LA supplementation significantly reduced IBA-1 expression of obese mice also at cortical level, and in particular in the PL [Z=2.3, p<0.05], OFC [Z=3.1, p<0.01], ACC [Z=2.1, p<0.05] and aIC [Z=2.2, p<0.05] (no main effect observed in the IL; KW, p>0.05) (**Figure 4**). Concerning the subcortical regions investigated where a significant increase of IBA-1 expression consequent to the rich-fat regime was observed, a beneficial effect of LA administration has been revealed in the DLS [Z=2.5, p<0.05], dBNST [Z=3.0, p<0.01], BLA [Z=2.2, p<0.05], LHb [Z=2.2, p<0.05] and VPPC [Z=2.5, p<0.05], as well as a tendency in the DMS [Z=1.9, p=0.05] and CeA [Z=1.8, p=0.07] (no main effect observed in the dHipp and VPM; KW, p>0.05) (**Figure 5**).

The effect of diet LA supplementation was also evaluated in non-obese animals. From the 19 regions investigated, IBA-1 expression was increased only in the CeA [Z=-2.4, p<0.05], PSTN [Z=-3.1, p<0.01] and STN [Z=-3.2, p<0.01] of STD-LA mice compared to control animals, and a tendency was observed in the aIC [Z=-1.9, p=0.06], LHb [Z=-1.7, p=0.08] and BLA [Z=-1.9, p=0.06]. No other effects were observed concerning LA administration in mice under a STD regime (all ps>0.1) (**Figure 3-5**).

### Correlations between behavioural scores and IBA-1 expression

Aiming at better understanding the relationship between microglial-related neuroinflammation and the emergence and installation of anxio-depressive-like symptomatology in HFD-induced obesity, a correlation study confronting IBA-1 fluorescence intensity and behavioural scores was carried out.

Among all the brain regions studied, five main anatomical actors, i.e. DMS, dHipp, LHb, VPM and STN, might explain a relationship between microgliosis and the behaviour observed in the LDB test. Thus, mice with higher fluorescence intensities related to IBA-1 in these brain regions consistently spent shorter periods in the transition zone, i.e. they show a decreased exploratory behaviour compared to those with fewer intensities [DMS, r=-0.426, p<0.05; dHipp, r=-0.594, p<0.001; LHb, r=-0.597, p<0.001; VPM, r=-0.695, p<0.0001; STN, r=-0.599, p<0.001], and visit less often the light compartment [dHipp, r=-0.372, p<0.05; LHb, r=-0.341, p=0.07; VPM, r=-0.436, p<0.05; STN, r=-0.315, p=0.10]. However, as illustrated in **Figure 6a-j**, the variation of the data set explained by these linear models (R^2^ values) does not reach 50% for the overall of individuals. Interestingly, if the pathological experimental group (HFD-VEH) is studied separately, the explained variation of the data set considerably increases [HFD-VEH, time in transition: dHipp, r=-0.759, p<0.05; LHb, r=-0.726, p<0.05; VPM, r=-0.894, p<0.01; STN, r=- 0.758, p<0.05; entries in light: DMS, r=-0.563, p=0.07; dHipp, r=-0.815, p<0.05; LHb, r=-0.790, p<0.05; VPM, r=-0.906, p<0.01; STN, r=-0.787, p<0.05], which is not the case for the other experimental groups [STD-VEH, STD-LA and HFD-LA, time in transition: DMS, r=-0.045, p=0.84; dHipp, r=-0.360, p=0.11; LHb, r=-0.405, p=0.07; VPM, r=-0.545, p<0.05; STN, r=-0.398, p=0.07; entries in light: DMS, r=-0.338, p=0.15; dHipp, r=-0.055, p=0.82; LHb, r=-0.022, p=0.92; VPM, r=-0.169, p=0.46; STN, r=-0.042, p=0.86]. These results therefore suggest that microglial-related response might account for the behaviour+-characterizing HFD-induced obese individuals in the LDB, but not for the same behaviour in non-obese individuals (STD-VEH and STD-LA), or obese individuals where neuroinflammation has been rescued (HFD-LA).

**Figure 6.**
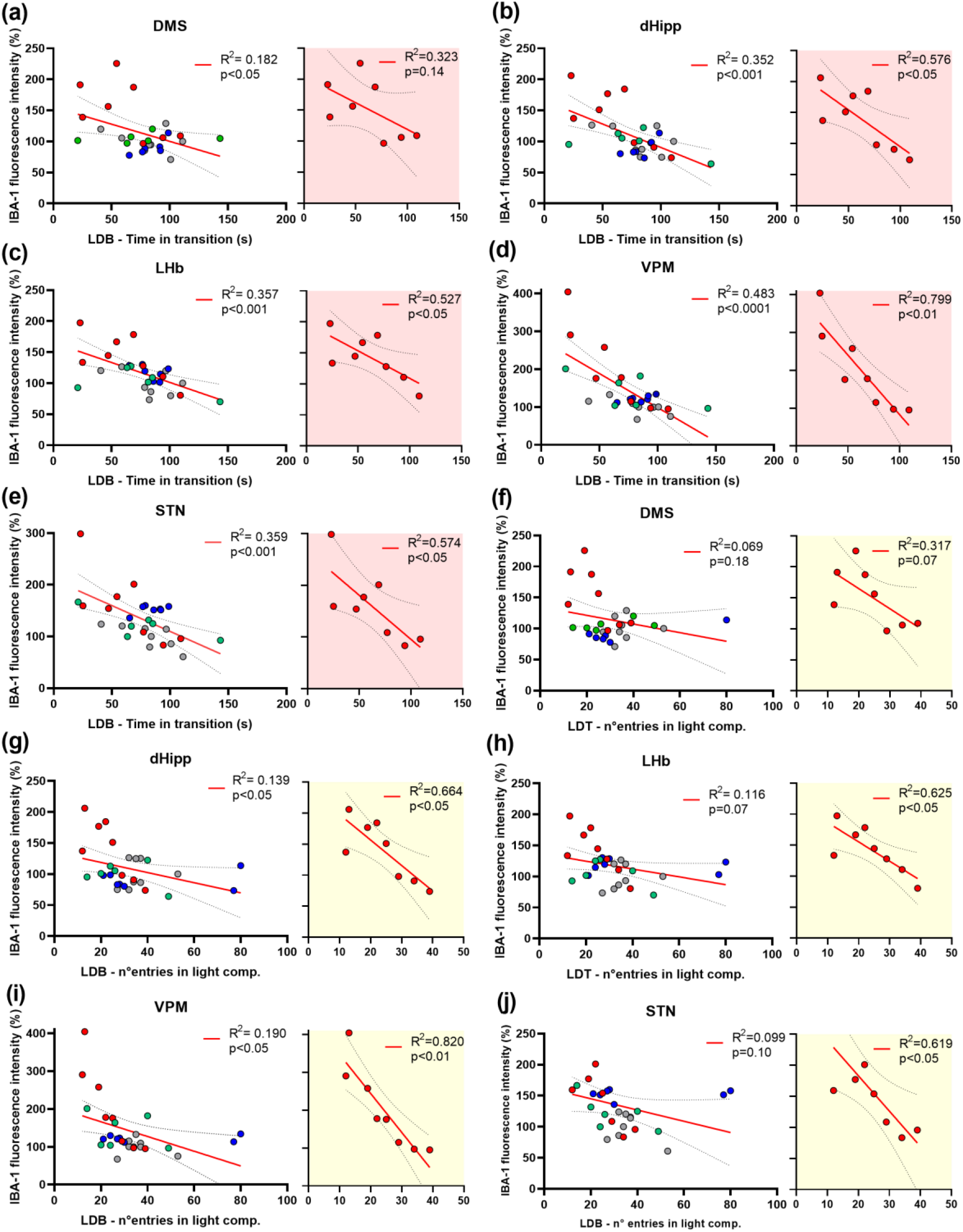
Relationship between behavioural scores in the light-dark box (LDB) test and IBA-1 relative expression in the studied brain regions. From the overall results, the correlations studies reveal five main anatomical actors (dorsomedial striatum -DMS, dorsal hippocampus -dHipp, lateral habenula - LHb, ventral posteromedial thalamic nucleus -VPM, and subthalamic nucleus of the hypothalamus - STN) that might explain 2 behavioural scores in the LDB test, i.e. time in the transition zone **(a-e**, white plotting area) and number of entries in the light compartment **(f-j**, white plotting area). However, the explained variation of the data set by the linear models is globally poor (<50%). Besides, when the pathological and non-treated condition is isolated (HFD-VEH), the explained variation of the linear models notably increase **(a-e**, time in transition: red plotting area; **f-j**, entries in light: yellow plotting area). The results suggest that, in pathological conditions, IBA-1 expression might account for behavioural variability in the LDB test in the highlighted regions. Colour code: grey for STD-VEH, blue for STD-LA, red for HFD-VEH and green for HFD-LA.

Concerning the scores calculated for the other behavioural tests, only three other significant correlations have been evidenced. For the total time of grooming in the ST, overall scores significantly correlate with IBA-1 intensity in the DMS [r=-0.401, p<0.05], but not for HFD-VEH if analysed separately [r=-0.512, p=0.20]. For the latency of approximation in the NSF task, overall scores significantly correlate with IBA-1 intensity in the PSTN [r=0.513, p<0.01], but not for HFD-VEH individuals separately [r=0.384, p=0.40]. However, NSF latencies significantly correlate with IBA-1 intensity in the PLH of the HFD-VEH experimental group [r=0.716, p<0.05]. No other significant correlations have been found between behavioural scores and IBA-1 fluorescence intensities in the brain regions studied [all ps>0.10].

## Discussion

Mechanisms underlying chronic central inflammation consequent to sustained consumption of saturated high-fat regimes remain uncertain. Furthermore, excess of fat intake and obesity enhance the individuals’ vulnerability to suffer from emotional disturbances [5,8,10–12,33]. In this frame, the here presented experimental investigation shows that saturated HFD-induced neuroinflammation extends beyond the hypothalamic region, with a significant impact also at cortical and subcortical levels. Neuroinflammation was concomitant with the emergence of an anxio-depressive-like symptomatology in obese mice, but further investigation is needed in order to identify a causal relationship. Most importantly, this experimental work confirms *in vivo* anti-neuroinflammatory properties of LA supplementation, and demonstrates its therapeutic potential at the chronic stage of central inflammation in a murine model of diet-induced obesity.

In line with previous studies [11,34–36], this investigation shows that sustained consumption of fatty diets leads to a significant increase of the body weight that, to a certain extent, is a consequence of an installed hyperphagia. Growing scientific evidence associates the behavioural changes observed in obese individuals concerning food intake (food choice and amount of consumed food), to reduced taste perception [37–39]. Therefore, the consummatory behaviour observed in the HFD experimental group may be explained as a consequence of a lipid-specific taste dysfunction resulting from reduced oral bud cell density and/or inhibited renewal. Furthermore, orosensory detection is known to be sensitive to systemic inflammation [40,41], which reinforces the idea of a positive-feedback loop that progressively aggravates the metabolic and emotional profile of individuals suffering from obesity.

Obesity-related emotional disturbances arise in a comorbid manner to metabolic and systemic impairments. Hence, the emergence of anxio-depressive-like phenotypes consequent to sustained consumption of saturated HFD has been proved in rodent models [10]. Consistently, the results of the here presented behavioural evaluation show that HFD-induced obese mice display typical anxio-depressive-like behaviours in the ST and the LDB tests. Compared to control individuals, the grooming time and the time invested exploring the light compartment is reduced in obese mice. However, the other parameters evaluated in the LDB, as well as the NSF task, did not succeed in revealing emotional disturbances. In the specific case of the NSF task, the unexpected results obtained during the evaluation suggest that the chosen anxio-depressive index might not be appropriate. The latency to first approximate food was supposed to sufficiently illustrate the HFD-induced emotional misbalance in obese mice [42], but literature also refers to the use of the latency to start eating as an index of motivational impairment in the NSF task [43]. Therefore, the lack of difference between NSF behavioural scores in obese vs non-obese individuals should be interpreted with caution. Finally, HFD-induced obesity in mice did not induce behavioural modifications in terms of repetitive compulsive-like behaviour in the NST. A specific behavioural output was not expected for obese individuals in the NST, but since repetitive behaviours have been often described in the literature in animal models of anxiety/depression [44], this evaluation was included in the experimental design in order to gain insight.

Although literature reports beneficial contributions of dietary supplementation on emotional disturbances consequent to sustained consumption of saturated high-fat regimes [13,45,46], neither anti-depressive nor anxiolytic effects of five weeks of LA supplementation were observed in the present investigation. In an interesting experimental approach, Demers and colleagues [13] demonstrated that when a saturated HFD-regime was supplemented with fish oil rich in polyunsaturated ω-3 fatty acids, the anxio-depressive-like symptomatology of obese mice was alleviated. Even though the treatment durations were equivalent, the protocols used to administrate fish oil (gavage) and LA (*ad libitum* drinking beverage) were different and may account, to a certain extent, to the differences observed between studies. However, while five weeks of fish oil administration attenuated hyperphagia, LA supplementation did not significantly change food intake in HFD-induced obese mice. Complementary metabolic analyses (e.g. glucose tolerance) in this context will help better understanding while five weeks of LA supplementation were not sufficient to improve the emotional state of obese mice.

Systemic and central inflammation are characteristic of obesity. Neuroinflammation consequent to sustained high-fat regimes has been well described in hypothalamic regions due to their key role on energy homeostasis and food intake regulation [for review ^47^]. However, the inflammatory profile of obesity extends to the entire CNS, as this study have demonstrated. Microgliosis, as a proxy of neuroinflammation (since microglia are the immune cells of the CNS) characterize hypothalamic, cortical and subcortical brain regions of obese mice, with few exceptions. Microglia activation consequent to high-fat regimes occurs notably early. Indeed, 1-to-4 days of HFD are sufficient to induce changes in this cell population [48,49], but literature alludes to the chronicity of neuroinflammation as a key feature for the emergence and installation of emotional disturbances. Based on previous observations and studies [31,50], this investigation evaluated the anti-neuroinflammatory properties of LA while supplementing a saturated HFD, as well as its impact on obesity-induced anxio-depressive-like behaviours. Five weeks of LA administration rescued microglia-related neuroinflammation in 11 of 13 brain regions investigated, and three tendencies were also observed (13 of 20 regions with significant higher IBA-1 related fluorescence intensities in HFD-VEH vs STD-VEH). Therefore, since LA exerts overall anti-inflammatory effects in the brain, an improvement of the emotional disturbances consequent to sustained HFD was expected. Unfortunately, the anxio-depressive-like phenotype of obese mice in the ST and the LDB test did not benefit from LA supplementation. One possible explanation for the lack of behavioural effect is the duration of the treatment. Evaluating the effect of LA supplementation on the rescue of microglia-related neuroinflammation at different time-points will certainly help understanding the kinetics of the underlying mechanisms.

Another interesting result of this investigation was the increased IBA-1 expression in the STN and PSTN of control mice after 5 weeks of LA supplementation. Both structures are known to participate in the regulation of food intake, and in particular the PSTN, which is involved in stopping feeding after a negative reward evaluation [51]. Besides, it has been shown that STN optogenetic manipulation modifies food intake, but its particular role in feeding behaviour in still unclear [52]. Thus, the isolated LA effect suggest that these hypothalamic structures are especially sensitive to the treatment and therefore, to dietary variations of fat intake. One possible explanation would be a different origin of the inflammatory response. In addition to an undercharacterized effect of central inflammation in terms of regional specificity and time course, the neural-glial dynamics of HFD-induced hypothalamic inflammation are not clearly understood. We hypothesized that STN and PSTN might act as high-sensitive sensors of fat variations within the hypothalamus, and enhance the inflammatory response through different synergistic effects of neural-originated and glia-originated inflammation. Therefore, it would be interesting to evaluate the implication of the specific inflammatory pathway of IKKβ signalling in these specific areas, given that its dysregulation has been linked to HFD-induced neuroinflammation and weight gain [53].

Finally, in an attempt to find plausible causal relationships between the behavioural scores evaluated and the severity of the central inflammation, a correlation study was carried out including all four experimental conditions. Significant correlations were found for five anatomical actors (four subcortical and one hypothalamic region) and two LDB behavioural scores (time in transition and entries in light compartment). Intriguingly, no HFD-induced microgliosis was evidenced in two of these structures (dHipp and VPM), and no significant emotional difference revealed for the indicated behavioural scores. However, IBA-1 fluorescence intensities calculated in the dHipp and VPM of obese mice show a wide dispersion, suggesting different neuroinflammatory profiles that may account for the lack of difference. In any case, while linear models did not succeed at explaining the overall variation of the data sets, they eventually better predicted microgliosis through behavioural scores in non-treated obese individuals. These results suggest that using microglia-specific IBA-1 expression to explore the relationship between neuroinflammation and emotional misbalance might require different mathematical approximations depending on the microglial activation state. Thus, other complementary measurements should be considered in order to better illustrate the severity of the central inflammation.

Together, these findings support dietary LA supplementation for alleviating neuroinflammation associated to overconsumption of fatty food and obesity. Nevertheless, further investigation is necessary to untangle the relationship between neuroinflammation and HFD-induced obesity in mice. Since neuropsychiatric disorders as anxiety and depression exacerbate the risk that obesity burdens on human health, finding successful therapeutic strategies is necessary. The results of this study are therefore relevant to understand the pathophysiological mechanisms underlying emotional-based behaviours in diet-induced obesity.

## Methods

### Animals

Forty C57BL/6JRj male mice (*Ets Janvier Labs*, Saint-Berthevin, France) aged of 8 weeks at the beginning of the experiments were used in this study. Mice were group housed and maintained under a normal 12-h light/dark cycle with controlled temperature (22±2 °C) and environmental humidity (55±10%). They had *ad libitum* access to food (standard diet –STD, or high-fat diet –HFD; see **Food regime** section for details) and drinking water, vehicle or LA supplementation/treatment (see **Treatment** section) during the entire duration of the experiment, and their weight was regularly monitored (mean weight ± SEM (g) at the beginning of the experiment, STD: 26.46±0.33; HFD: 26.74±0.35).

The procedures were all conducted in accordance with the European Community Council Directive of September 22^nd^, 2010 (2010/63/UE), and following the standards of the Ethical Committee in Animal Experimentation from Besançon (CEBEA-58; C-25-056-10; MESRI-APAFiS#28240).

### Food regime

After a 4 days-period of habituation to the animal facilities, mice were randomly divided into two groups of 20 individuals each, and were fed with either a SD (KlibaNafag3430PMS10, *Serlab*, CH-4303 Kaiserau, Germany) or a HFD (A03 food powder, *SAFE*, Augy, France; cholesterol C8667, *Sigma-Aldrich*, France; palm oil, *Vigean*, France; as previously described [34]) for the entire duration of the study (17 weeks). Global house food intake was measured in parallel to body weight monitoring.

### Treatment

After 12 weeks of differential regime, mice were randomly divided in 4 groups of 10 individuals each: SD and HFD mice treated with vehicle (VEH; 0.3% Arabic gum solution; G9752, *Sigma-Aldrich*, France), and SD and HFD mice treated with a LA solution (0.2% LA in VEH; L1376, *Sigma-Aldrich*, France). The drinking solutions were prepared every 2-3 days, and their drinking intake was systematically measured.

### Behavioural characterization

The behavioural evaluation of the animals started after 5 weeks of VEH/LA administration. Tests were conducted during the light phase of the circadian cycle (starting at 8 a.m., inactive phase) in an adapted behavioural room.

#### Nestlet shredding test (NST)

Repetitive, compulsive-like behaviours were assayed using the NST, which can robustly measure the effects of different treatments on the natural and spontaneous nestlet shredding behaviour of mice. For that, animals were individually placed into a cage with litter and a preweighted nestlet, and left undisturbed for 30 min. After the test completion, mice were replaced in their home cages and the remaining intact nestlet material was removed and allowed to dry overnight. The unshredded material was then weighted in order to calculate the percentage of the total nestlet shredded, which served as index of compulsive-like behaviour [54].

#### Splash test (ST)

Depressive-like behaviour was evaluated using the ST, based on the internal drive of rodents to groom themselves. Briefly, each animal was individually placed for 5 min in an individual home cage with litter after the dorsal surface of its body was sprayed with a 15% sucrose solution. A longer latency to the first grooming period and an increase in the total time of grooming are considered indicators of a depressive-like phenotype [55].

#### Novelty suppressed feeding (NSF) task

The inhibition of feeding produced by a novel environment or hyponeophagia can be measured in rodents using this task. The NSF task informs about alterations of the internal state of individuals by appealing to the appetitive component of incentive motivation [43], and is sensitive to pharmacological treatments targeting anxio-depressive symptomatology [56,57].

Animals were food-deprived for 12 hours before being individually placed in an opaque cylindrical open-field apparatus (height: 60 cm; diameter: 47 cm), in the centre of which animals could reach two grain-based pellets located on a white piece of filter paper (diameter: 10 cm). Similarly to previous studies [58,59], the behaviour of animals was recorded, and their latency the first approach to food was used as index of anxio-depressive-like behaviour. The test lasted a maximum time of 10 min.

#### Light-dark box (LDB) test

The light-dark box (LDB) test allows evaluating unconditioned anxious responses in rodents, by exposing the animals to the internal conflict of choosing between remaining in a dark compartment (perceived as safe; width: 40 cm; high: 14 cm; 20-30 lux) or to explore a bright and open compartment (i.e. to expose themselves in a novel environment; width: 40; high: 26; 120-130 lux). The test lasted 10 min, during which the animals’ behaviour was recorded and sampled using an automatic videotracking system (*Viewpoint Behavior Technology*, Lyon, France). The parameters evaluated were: the time spent and the entries in the light compartment, the time spent in the transition zone, and the total distance travelled in the box. Entries were considered when the four legs of the animals were placed in one of the compartments. When the body of the animal was placed between the two compartments, it was considered that the animal was in the transition zone (adapted from ^35^).

### Animals sacrifice and tissue sampling

Twenty-four hours after the completion of the behavioural assessment, animals were sacrificed for evaluation of the neuroinflammation by IBA-1 immunofluorescence. The procedures were similar to those previously described by our team [58,59]. Briefly, after an intraperitoneal injection of pentobarbital (55 mg/kg, Exagon®, *Med’Vet*, France), mice were transcardially perfused with 0.9% NaCl, followed by icecold 4% paraformaldehyde fixative (PFA, *Roth®*, Karlsruhe, Germany) in 0.1 M phosphate buffer (PB, pH 7.4). Brains were extracted and postfixed overnight in the same fixative at 4 °C and cryoprotected by immersion in a 15% sucrose solution (D(+)-Saccharose, *Roth®*, Karlsruhe, Germany) in a 0.1 M PB 24 h at 4 °C. After freezing by immersion in isopentane (2-methylbutane, *Roth®*, Karlsruhe, Germany), brains were cut in coronal 30-µm-thick serial sections and stored at −40 °C in a cryoprotector solution (1:1:2 glycerol/ethylene glycol/PB; *Roth®*, Karlsruhe, Germany).

### Immunofluorescence

Six-to-eight individuals of each experimental group were randomly selected for assessing neuroinflammation by the proxy of the microglial response. For that, IBA-1 expression was targeted in a selection of hypothalamic, cortical and subcortical brain regions directly or indirectly implicated in the homeostatic and/or hedonic regulation of food intake, and in the emergence of anxio-depressive-like symptomatology. The selected tissue sections were: prelimbic (PL), infralimbic (IL) and orbitofrontal (OFC) cortices at (1.94–1.70) anterior to Bregma (aB); medial anterior cingulate cortex (ACC), dorsolateral (DLS) and dorsomedial (DMS) striatum at (1.10–0.98) aB; anterior insular cortex (aIC), paraventricular nucleus of the hypothalamus (PVN) and dorsal region of the bed nucleus of the stria terminalis (dBNST) at (0.26–0.02) aB; amygdala (central -CeA and basolateral -BLA) and peduncular part of the lateral hypothalamus (PLH) at (1.06–1.34) posterior to Bregma (pB); dorsal hippocampus (dHipp), lateral habenula (LHb) and ventromedial hypothalamus (VMH) at (1.70–1.94) pB; ventral posteromedial thalamic nucleus (VPM), parvicellular part of the ventral posterior thalamic nucleus (VPPC), lateral hypothalamus (LH), subthalamic nucleus (STN) and parasubthalamic nucleus (PSTN) at (2.06–2.30) pB, according to Franklin and Praxinos [60]. After washing, floating sections were incubated with the primary antibody (1:2000; ab178846, rabbit anti-IBA1, *Abcam*) during 24 h at 4 °C. Tissues were then incubated with the secondary antibody (1:1000; A10520, goat anti-rabbit IgG Cyanine 3, *Invitrogen*) during 2 h at room temperature. Finally, sections were washed, mounted on gelatincoated slides and coverslipped with mounting medium (40% PB, 60% glycerol; *Roth®*, Karlsruhe, Germany). Photomicrographs of brain structures were acquired using 10x objectives of an Olympus microscope B x 51, equipped with a camera Olympus DP50. *ImageJ* software was used to quantify IBA1 related fluorescence over the regions of interest. Final samples sizes were: PL n=30, IL n=30, OFC n=30, medial aCC n=28, DLS n=28, DMS n=28, aIC n=29, PVN n=29, dBNST n=29, CeA n=29, BLA n=29, PLH n=29, dHipp n=29, LHb n=29, VMH n=29, VPM n=29, VPPC, n=29, LH n=29, STN n=29 and PSTN n=27.

### Data and statistical analyses

The results are presented as means ± SEM. The statistical analyses were conducted using STATISTICA 10 (*Statsoft*, Palo Alto, United States), and the figures were designed using GraphPad Prism 9 software (*GraphPad Inc.*, San Diego, United States). Assumptions for parametric analysis were systematically verified using Shapiro-Wilk and Levene’s tests to respectively study the normality of distribution and the homogeneity of variance of the data sets. The progression of the body weight, of the food intake and of the drinking beverage intake were evaluated with a repeated measures ANOVA (RMA) design including 2 (STD and HFD) or 4 (STD-VEH, STD-LA, HFD-VEH and HFD-LA) groups (between-subject factor: regime or treatment) and 12 or 5 measurements (within-subject factor: time). When the nature of the data sets did not meet assumptions for parametrical analysis (behavioural and immunofluorescence scores), Kruskal-Wallis (KW) and Mann-Whitney U (MWU) tests were used. Gehan-Breslow Wilcoxon (GBW) tests were used to compare NSF score curves and the relationship between behavioural scores and IBA-1 expression in the studied brain structures were analysed using Pearson correlations. For all analyses, the significance level was set up at p<0.05, and analyses with p≤0.1 are interpreted as trends.

## Acknowledgements

The authors thank the *Burgundy -Franche-Comté Region* for awarding the project “*Tasty Lipids*” in the category “*Envergure*”, which helped realizing the present research work. The authors also thank the *Burgundy -Franche-Comté Region* for granting a post-doc scholarship to one of the authors (LC). Finally, the authors thank the Animal Facilities of Besançon for the technical support and PhD Pierre-Yves Risold (LINC) for his helpful insight during the immunofluorescence experiments.

## Author Contributions

Conceptualization of the experimental design: LC, NK, VVW. Methodology, investigation and data acquisition: LC, LJ, SD, BR, CH, AH. Data analysis and interpretation: LC, LJ and VVW. Drafting of the original manuscript: LC, LJ, EH and VVW. Supervision: LC, VVW. All authors critically revised the work and approved the version to be published.

## Data availability Statement

The data presented in this study are available on request from the corresponding author.

